# Reference BioImaging to assess the phenotypic trait diversity of bryophytes within the family Scapaniaceae

**DOI:** 10.1101/2022.07.29.501959

**Authors:** Kristian Peters, Birgitta König-Ries

## Abstract

Macro- and microscopic images of organisms are pivotal in biodiversity research. Despite that bioimages have manifold applications such as for assessing the diversity of form and function, FAIR bioimaging data in the context of biodiversity are still very scarce, especially for difficult taxonomic groups such as bryophytes. Here, we present a high-quality reference dataset containing macroscopic and bright-field microscopic images documenting various phenotypic attributes of the species belonging to the family of Scapaniaceae occurring in Europe. To encourage data reuse in biodiversity and adjacent research areas, we annotated the imaging data with machine-actionable meta-data using community-accepted semantics. Furthermore, raw imaging data are retained and any contextual image processing like multi-focus image fusion and stitching were documented to foster good scientific practices through source tracking and provenance. The information contained in the raw images are also of particular interest for machine learning and image segmentation used in bioinformatics and computational ecology. We expect that this richly annotated reference dataset will encourage future studies to follow our principles.

## Background & Summary

In biodiversity, organisms are studied with the aim to record their diversity at the genetic, metabolic, physiological, morphological, or the ecosystem level. Despite the fact that bioimaging techniques such as macro- and microscopy are prominently used, FAIR bioimaging data in the context of biodiversity are still very scarce^1–3^. This is especially the case for taxonomically difficult and underrepresented groups such as bryophytes. Currently, there are approx. 24’000 species of bryophytes known to science^4^. Unlike vascular plants, bryophytes lack well-differentiated organs that protect them from environmental exposures and pathogens. As a result, phenotypes are often cryptic and difficult to identify visually as they have developed unique specialised metabolisms and cell structures such as oil bodies^5,6^.

The highly diverse family of Scapaniaceae contains 48 taxa in Europe^7^ and is an ecologically important group regarding environmental adaptations^8^, the biochemistry of terpenoid natural products and other chemical structures^9–12^, the metabolism of pollutants and heavy metals^13,14^, and phylogenetics^15–18^. Generally, there is a considerable lack of described traits in bryophytes^19^ and especially in liverworts such as Scapaniaceae, phenotypic traits to assess the diversity of form and function are understudied^20^.

Bioimaging data in the field of biodiversity is of high relevance as they allow to assess the phenotypic diversity through an analysis and assessment of images^2,21,22^. In the form of measurable phenotypic traits, biological images are the groundwork of many ecological studies^3,20,21,23,24^. Phenotypisation through recording images of anatomical and morphological attributes allows qualitative and quantitative measurements of molecular structures relating to genetics, molecular pathways and biotechnology^25–27^. Bioimages have also gained a lot of interest in citizen sciences and in the digitization of natural history collections and digital herbaria^28,29^. Furthermore, meta-synthesis methods, which synthesise disparate data sources spanning published case studies, have great potential to reveal context-dependencies within bioimaging research data^30^.

Macroscopy and microscopy are characterized by physical constrains resulting in diffraction and shallow depth of field^22,31,32^. From a technical perspective, our data employs two major methods to significantly extend the depth of field and increase the resolution of the composite images: image stitching (combining several images relative to the x- and y-axes of the visible accommodation to form an image with a larger frame)^33,34^ and multi-focus image fusion (merging multiple images at different focal planes of the z-axis in such a way that only regions in focus will contribute to the resulting image)^35,36^. In this regard, the raw data also allows to be reused for combining image fusion with computational super-resolution^22,37–40^.

Cloud infrastructural resources are able to execute computational workflows that combine data with computational analysis tools at a large scale^41,42^. However, there is still a considerable lack of data containing machine-actionable meta-data^1,3,43–46^. To foster provenance, any raw and segmented image in this data set has been associated with a rich set of contextual and expressive meta-data^47^, documenting the phenotypic attributes, and recording any digital image processing (i.e., increasing contrast, brightness, image fusion). The meta-data has been formatted with community-accepted semantics that allow for machine-actionable data-mining and to create scientific workflow modules that produce segmented composite images automatically by reusing the instructions contained in our meta-data^20,42,45,46,48^.

In this Data Descriptor, we present the principles to generate reference images from raw microscopic bioimaging data and show how individual images are associated with technical and expressive meta-data. Despite that we were able to associate our images with a rich set of meta-data, we ascertain that there is still a lack of usable ontological terms and schemas in bioimaging with regard to documenting image processing and associating individual images with phenotypic attributes^3,45^. Our high-resolution images allow for large prints and zooming into images to obtain critical details, which is particularly important for species identification and for computational image analyses, computer-assisted species recognition and identification^49,50^. Despite that we have deposited the data in the two specialised imaging repositories BioStudies (containing raw and pre-processed images which enable rapid use in, i.e., machine learning approaches in computational ecology) and Imaging Data Resource (containing pre-processed and fully segmented images to be rapidly reused by ecologists), we ascertain that there is still the need for connecting macro- and microscopic bioimaging data to biodiversity platforms^3,51^ such as iDigBio^52^ and GBIF^53^, or even the citizen scientists community-effort iNaturalist^54^. Our reference data framework facilitates the further integration of bioimaging data into other research disciplines^55^ and, thus, we want to inspire future data reuse and meta-synthesis in the fields of biodiversity and computational ecology.

## Methods

Representative voucher specimens were received from different herbaria. Table 1 lists all used voucher specimen and freshly collected samples. Fresh samples of *Diplophyllum taxifolium, Scapania cuspiduligera, Scapania gymnostomophila* and *Scapania subalpina* were additionally collected at various sites, put into envelopes on-site, identified and photographed afterwards. Information regarding the date, site (including geographical coordinates), habitat, substrate and other further information were collected.

**Table 1:**
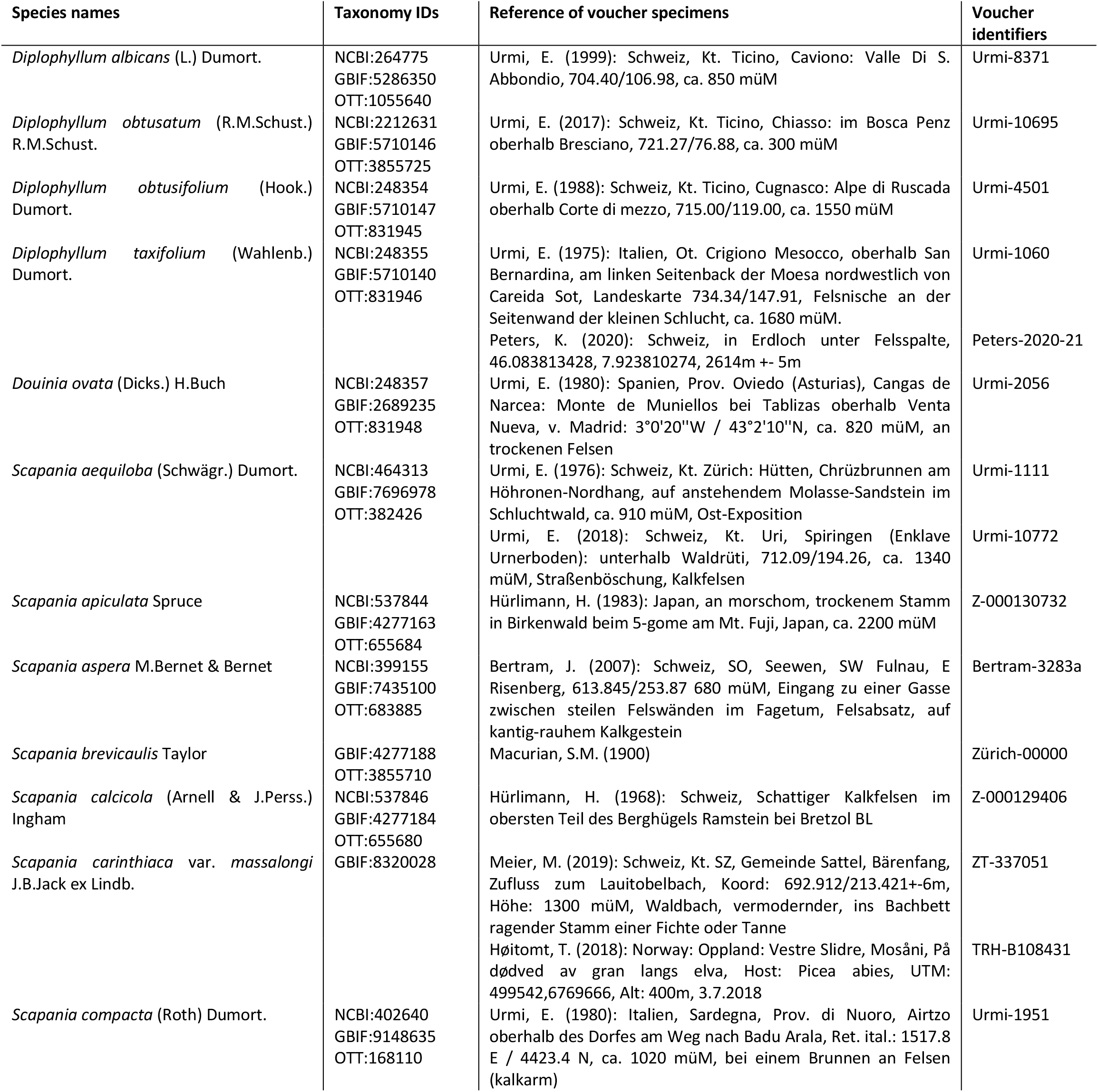
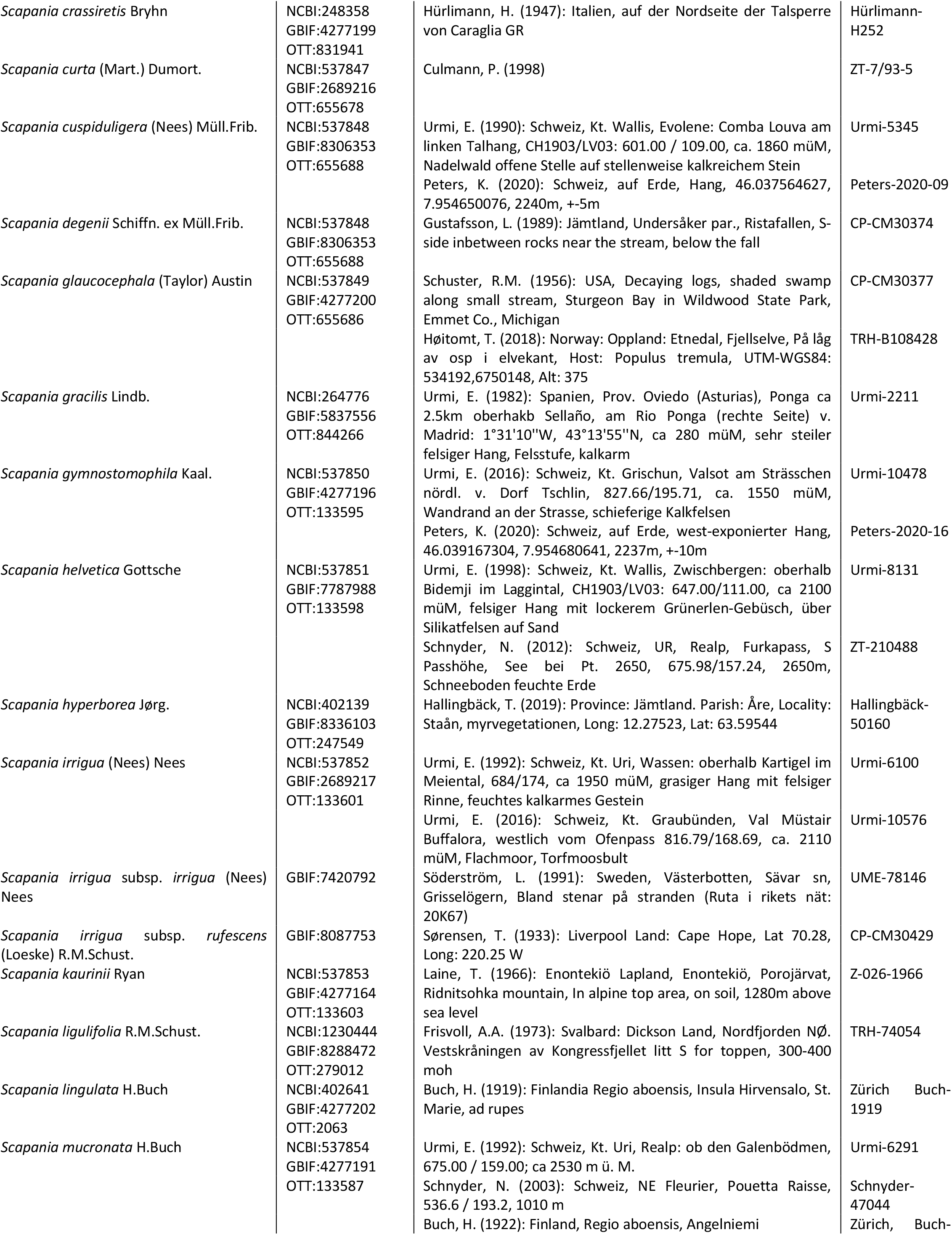
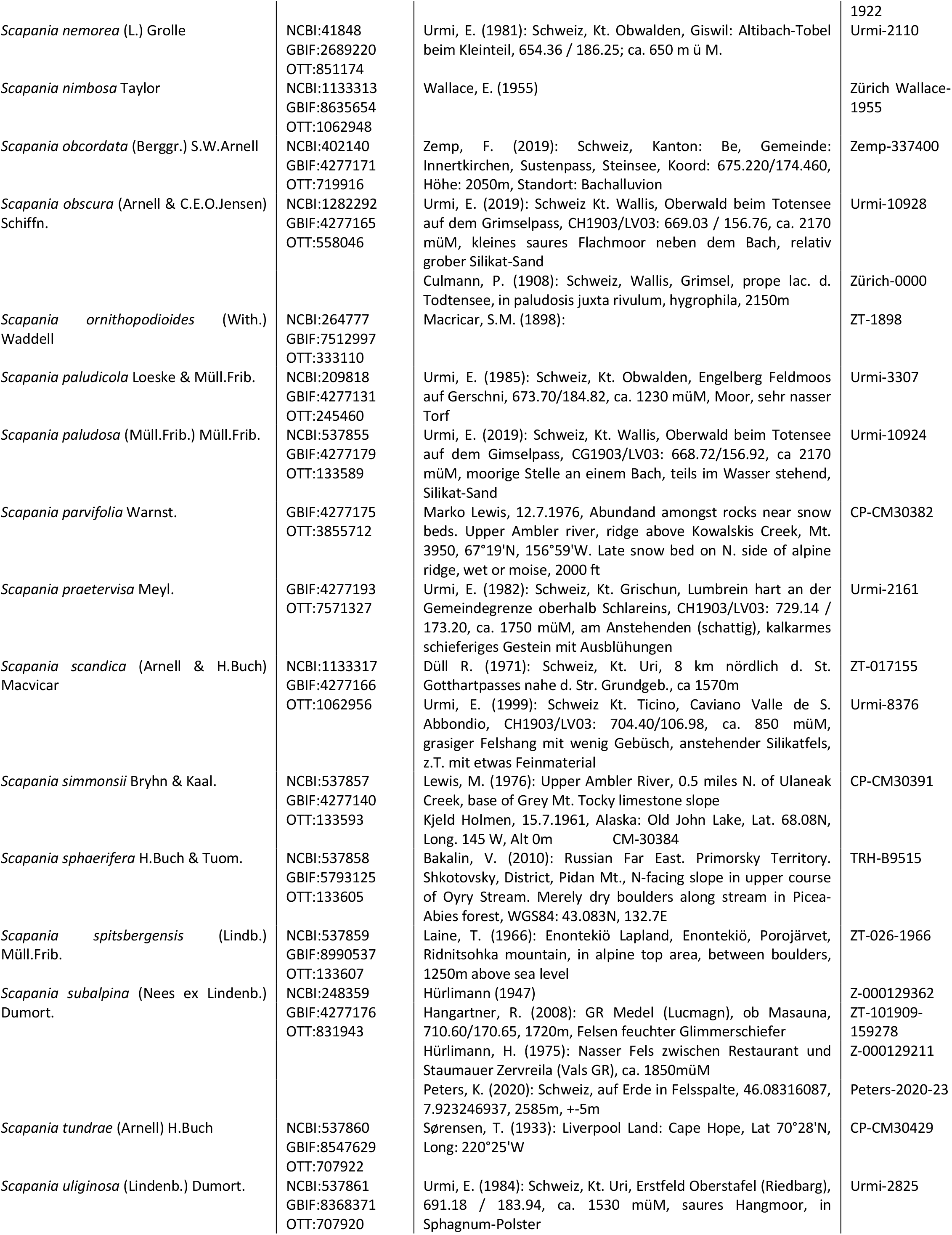
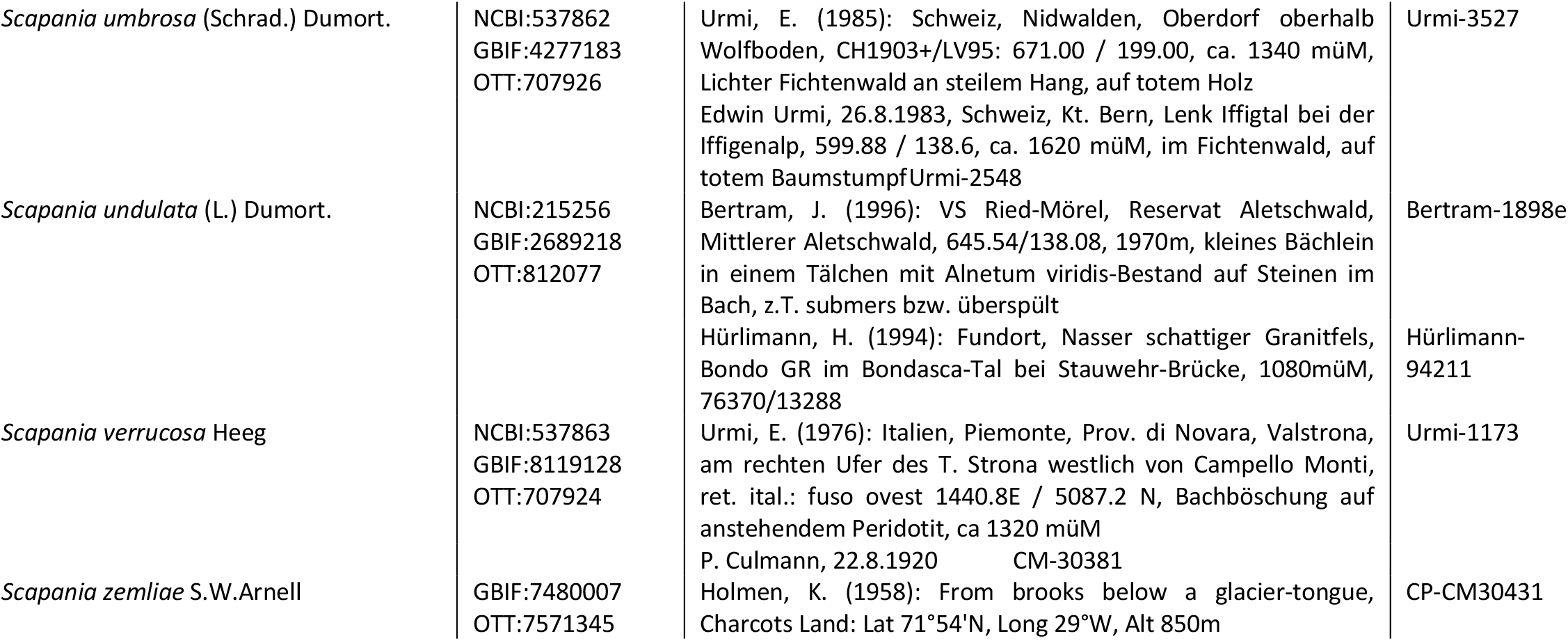
List of voucher specimens and fresh samples used in this study. All voucher specimens have been investigated. The columns list the taxonomic species identifiers (*NCBI, GBIF*, or *Open Tree of Life* identifiers, if available), the text on the specimen sleeves (collector, date and text on the envelopes) and the voucher specimen identifiers (the first letters either indicate the *Index Herbariorum* institution code^56^ if available or the name of private collection where the specimens were stored).

For microscopy, a Zeiss Axio Scope.A1 HAL 100/HBO, 6x HD/DIC, M27, 10x/23 microscope with an achromatic-aplanatic 0.9 H D Ph DIC condenser was used with the objectives EC Plan Neofluar 2.5x/0.075 M27 (a=8.8mm), Plan-Apochromat 5x/0.16 M27 (a=12.1mm), Plan-Apochromat 10x/0.45 M27 (a=2.1mm), Plan-Apochromat 20x/0.8 M27 (a=0.55mm), and Plan-Apochromat 40x/0.95 Korr M27 (a=0.25mm) using the EC PN and the Fluar 40x/1.30 III and PA 40x/0.95 III filters for DIC. The conversion filter CB3 and the interference filter wideband green were used to improve digital reproduction of colors. For macroscopy and for preparing microscopy slides, a binocular microscope Zeiss Stemi 2000c was used (apochromatic Greenough system with a stereo angle of 11° and 100/100 switchover of camera and ocular viewing). For stand-alone macroscopic images, the objectives Canon MP-E 65mm 2.8 1-5x macro and Laowa 25mm 2.5-5.0x ultra-macro for Canon EF and the Canon EF-RF adapter.

To acquire digital images, a full-frame, high-resolution camera (Canon EOS RP, 26 megapixel) was used and adapted to the microscopes using binocular phototubes with sliding prism 30°/23 (Axio Scope.A1) and 100:0/0:100 reversed image (Stemi 2000c) using 60-T2 camera adapter for Canon EOS and Canon EF-RF adapter. The objectives Canon MP-E and Laowa 25mm were adapted directly through the Canon EF-RF adapter.

To construct images with extended depth of field using computational methods, multiple images were recorded at different focal planes. This “focus stacking” approach was automatized for macroscopy by attaching the camera to a Cognisys StackShot macro rail fixed on a Novoflex macro stand, and for microscopy by adapting a Cognisys StackShot motor to the fine adjustment of the microscope using two cogged wheels, one small wheel (1 cm diameter) adapted on the motor and one large wheel (8.5 cm diameter) on the fine adjustment of the microscope. The two cogged wheels were coupled with a toothed belt to obtain very fine step increments of the stepping motor for high magnifications. A Cognisys StackShot controller was used to control the amount and distance of the stepping motor with the following controller settings: Dist/Rev: 3200 stp, Backlash: 0 steps, # pics: 1, Tsettle: 100.0 ms, Toff: 450.0 ms, Auto Return: yes, Speed: 3000 st/sec, Tlapse: off, Tpulse: 800.0 ms, Tramp: 100 ms, Units: steps, Torque: 6, Hi Precision: Off, LCD Backlight: 10, Mode: Auto-Step using between 25 steps (magnification 1x) and 50 steps (magnification 25x) and 100 steps (magnification 400x) (number of steps depending on aperture settings and effective magnification).

Raw images were recorded in CR3-format and pre-processed with Adobe Camera RAW. Non-destructive image processing such as corrections of the field curvature, removal of chromatic aberration, increase of contrast and brightness were performed in Adobe Camera RAW. Images were then exported to TIFF-format and any image prcoessing steps were recorded in individual Adobe XMP-files.

Multi-focus image fusion was performed on the individual images in the z-stacks using the software Helicon Focus 7.7.5 and by choosing the algorithms *depth map* and *pyramid* with different settings of radius (4, 8, 16, 24) and smoothing (2, 4). The best composite image was chosen manually and retained. When composite images contained specimen that were larger than the frame, several images were stitched together using the *panorama stitching* function in the software Affinity Photo 1.10.1.

Images were manually segmented and interfering background removed using the *flood select, brush selection* and *freehand selection* tools in the software Affinity Photo. In order to determine the scale, a stage micrometre was photographed separately with any of the objectives and microscope combinations. The scale was calculated per pixel for each combination (File scale_bar_distances.csv in ^57^) and scale bars were put post-hoc onto the segmented images using the Python script scale_bar.py^57^.

Meta-data including species name, taxonomic rank information (NCBI-Taxon, GBIF and OTT taxonomy identifiers), voucher specimen id, image acquisition date, an object description including the name of the captured phenotypic attribute(s), the used objective, microscope, and magnification were associated with any raw image based on unique respective file names. Individual file names (variable *file list*), name within an image focus stack (variable *stack name*) and name within an image stitching stack (variable *stitch name*) were recorded additionally to facilitate subsequent automized image processing in workflows. A Python script was created to put individual images as part of image stacks into directories (File create_image_stacks.py in ^58^). The Python script parses the *Label* tag in the XMP-files. Any meta-data regarding image enhancement and non-destructive image processing were extracted from XMP-files using a simple Python script (File xmp_stack_to_tsv.py in ^58^). The meta-data was saved in individual TSV-files and merged using a helper Python script (File tsv_merge.py in ^58^). Table 2 lists all fields which were extracted from the XMP.

**Table 2:**
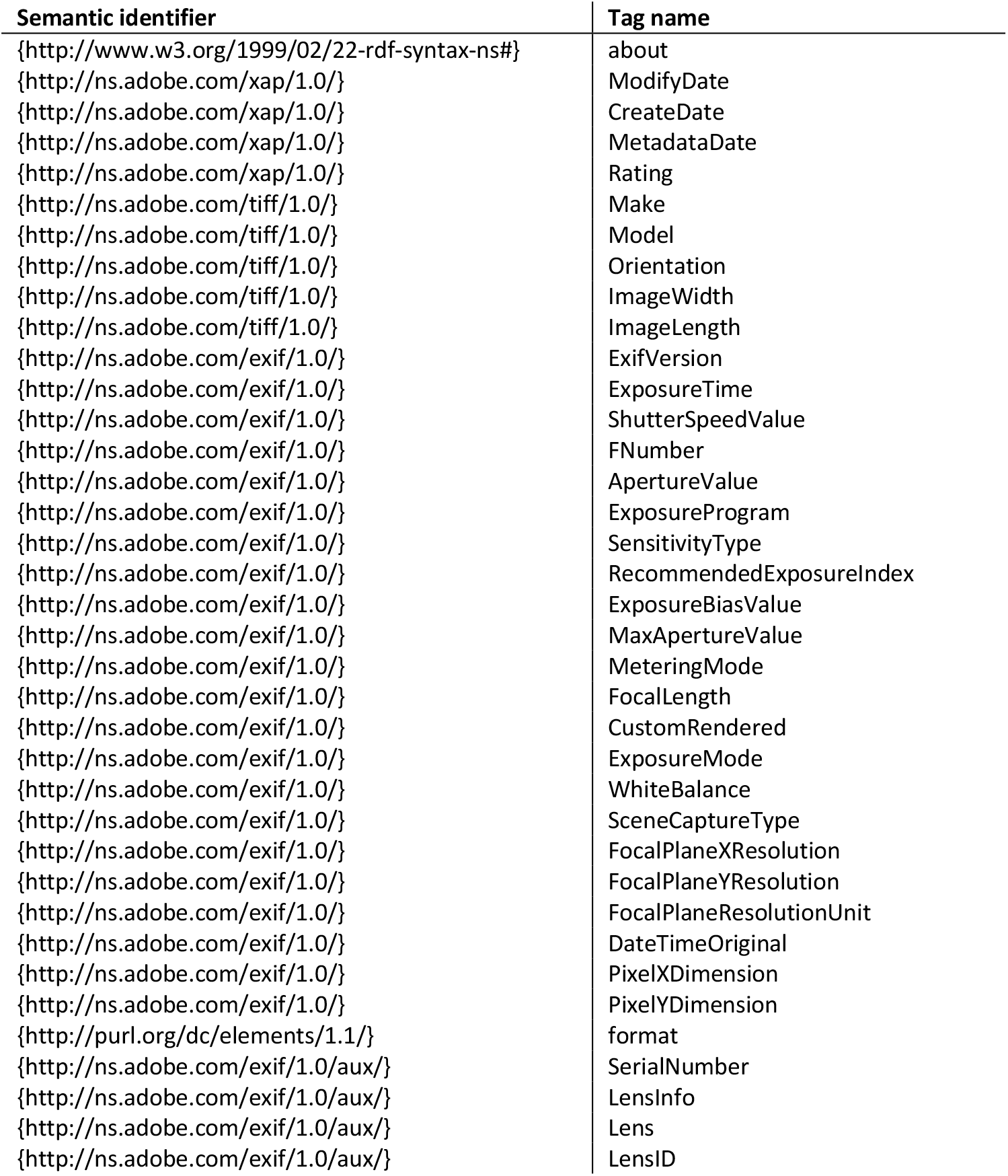
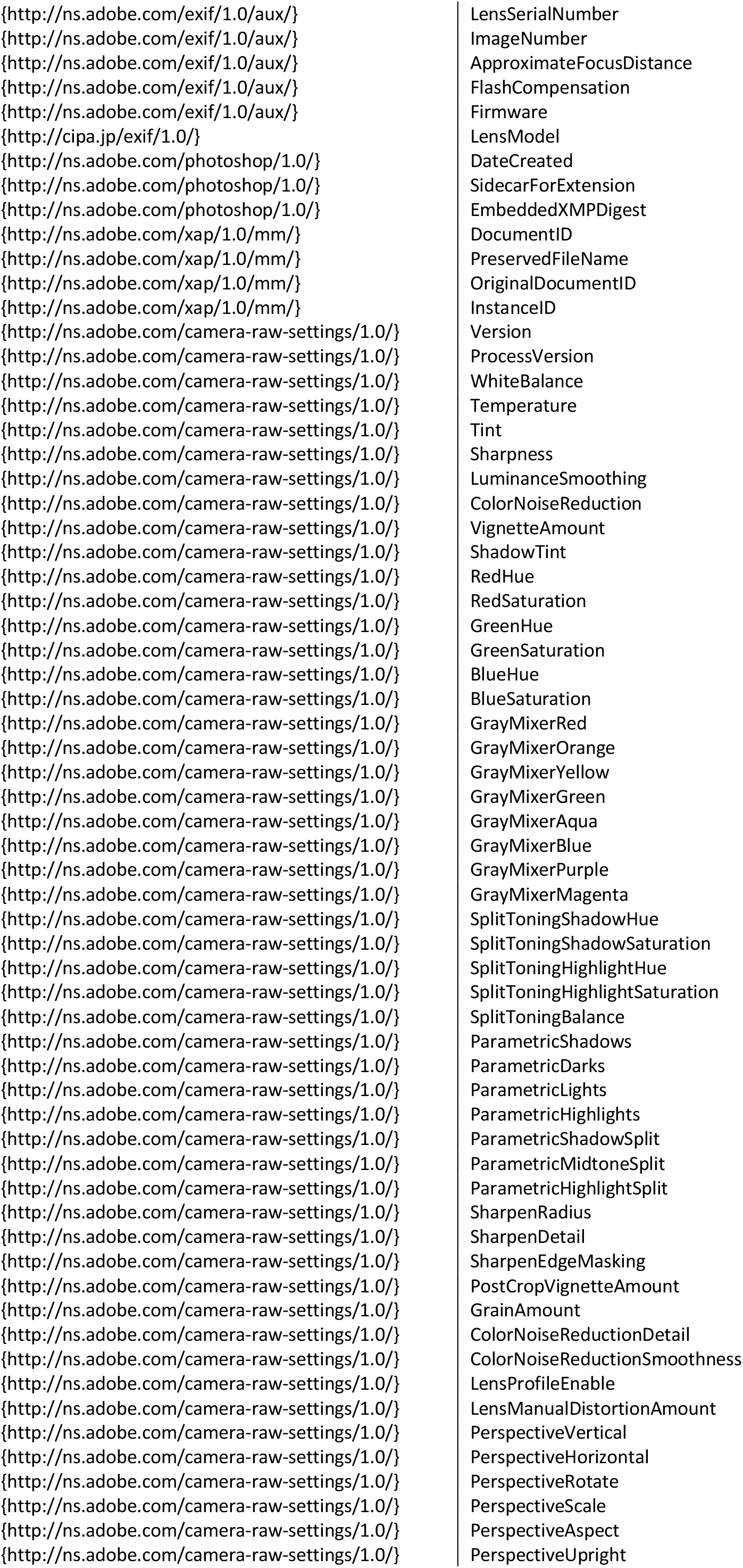
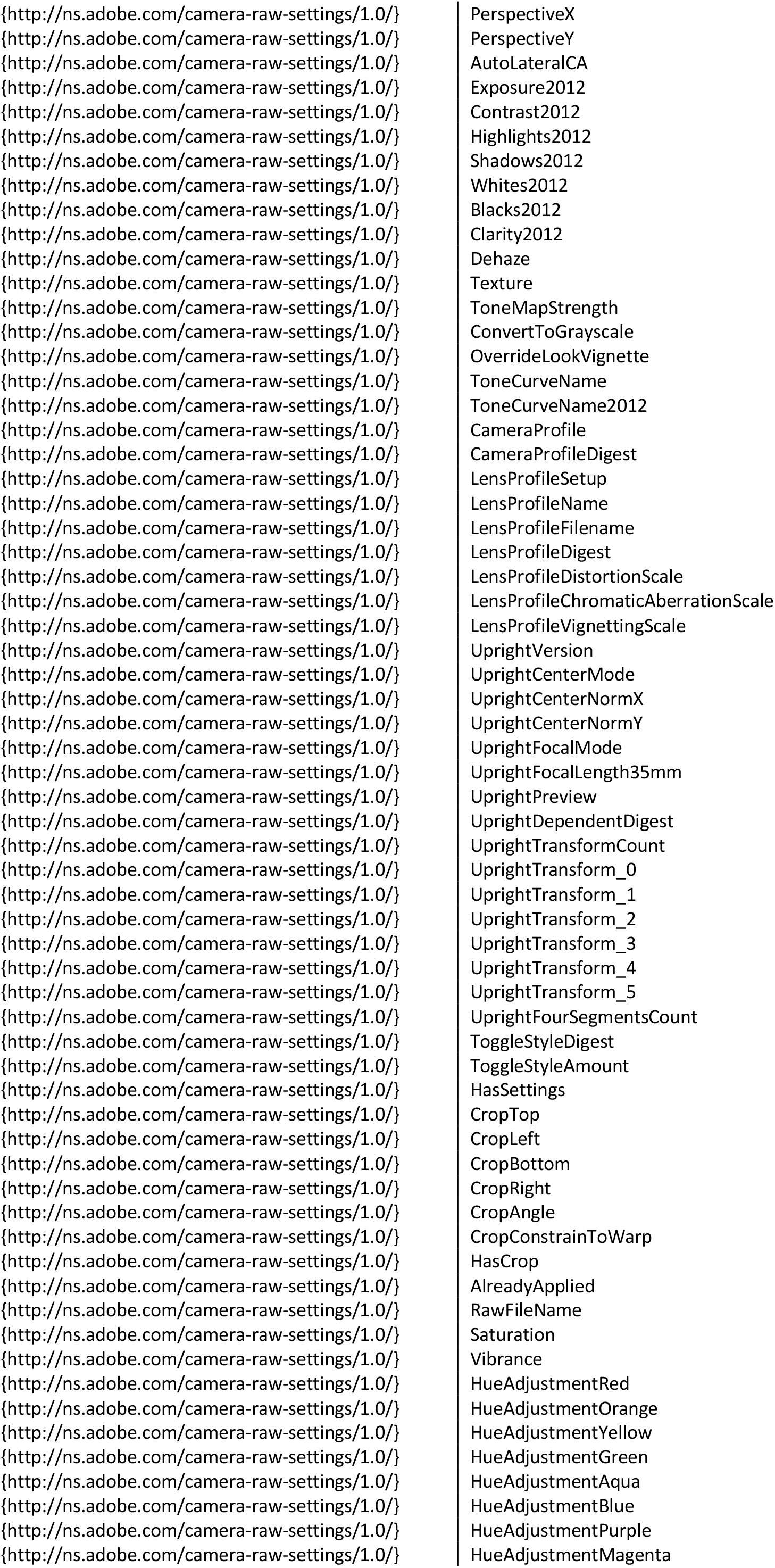
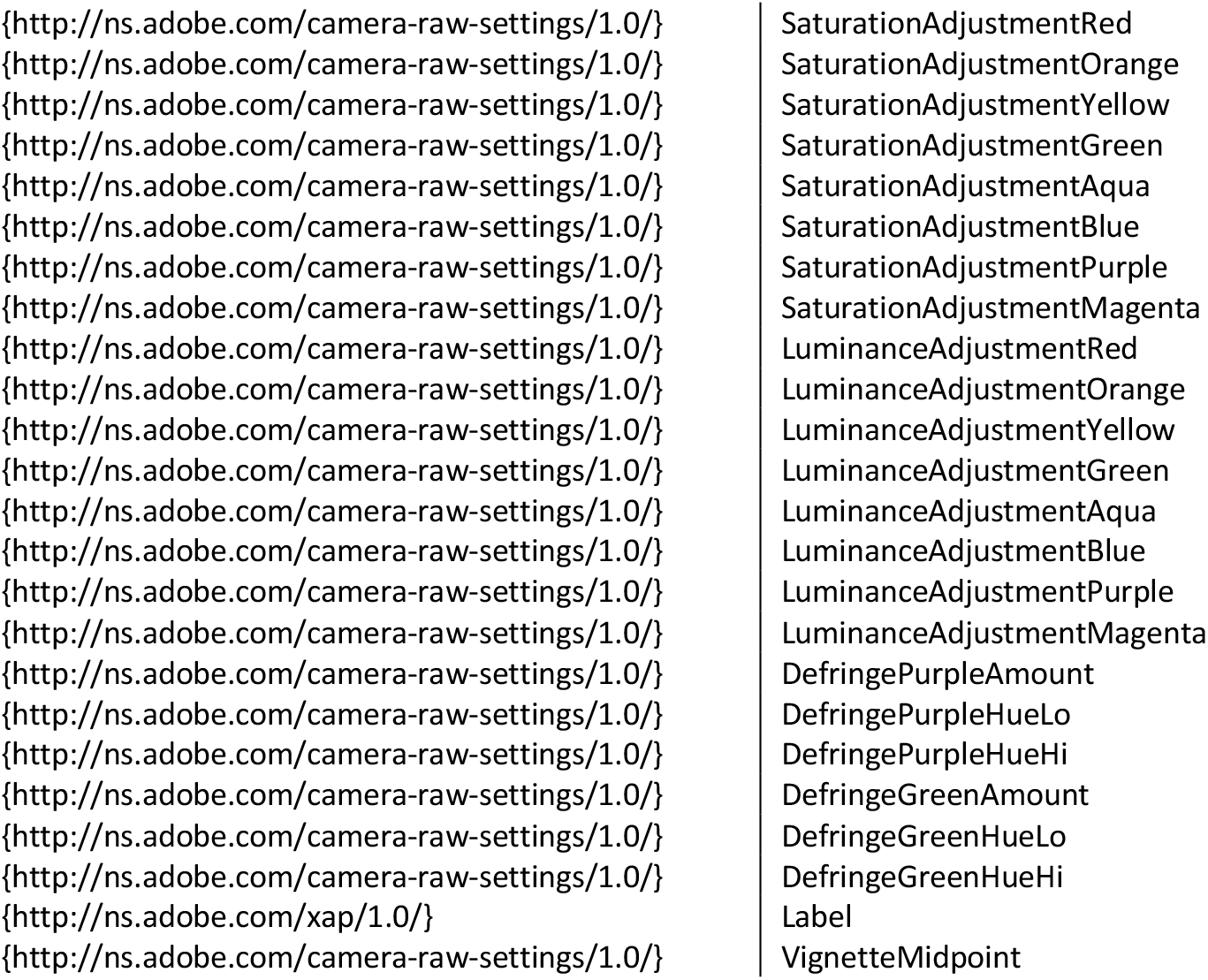
List of semantic identifiers used to annotate the image-enhancement performed in Adobe Camera-RAW.

Raw camera and pre-processed imaging data in CR3 and TIFF format, respectively, were uploaded to BioStudies using the command line IBM Aspera software tool *ascp* version 3.8.1.161274 to ensure that data has been transmitted without errors. Sparse file check summing was enabled to ensure integrity of files during transfer (parameter *-k 2*). The raw bioimaging data is available under the BioStudies identifier S-BIAD188.

Pre-processed images were converted to the Bio-Formats OME-TIFF format^59^ by creating intermediate ZARR-pyramid tiles using the bioformats2raw converter version 0.4.0 and then using the raw2ometiff version 0.3.0 software tool to create the final pyramid images. In order to improve data reuse and to enable linking bioimaging data to ecological data repositories, individual fully segmented and processed images were associated with standardised geolocation information. Swiss Topo CH1903/LV03 coordinates were converted to WGS84 using Swisstopo-WGS84-LV03^60^. The processed images were further associated with the meta-data information listed in Table 3. A helper script was implemented in R to facilitate the generation of TSV tables for data upload to the Image Data Resource (IDR) repository (_tsv_res_2_idr.r in ^61^). Processed images and the meta-data aggregated in a TSV table were uploaded to IDR using the software Globus Connect Personal 3.1.6. The dataset is available under the identifier idr0134.

**Table 3:** List of semantic terms used to annotate the segmented images.

| Semantic identifier | Name |
| --- | --- |
| {http://edamontology.org/data_1060} | File base name |
| {http://rs.tdwg.org/dwc/terms/acceptedScientificName} | Species name |
| {http://rs.tdwg.org/dwc/terms/TaxonID} | TaxonID |
| {http://rs.tdwg.org/dwc/terms/measurementMethod} | Measurement Method |
| {http://www.openmicroscopy.org/rdf/2016-06/ome_core/Instrument} | Microscope |
| {http://www.openmicroscopy.org/rdf/2016-06/ome_core/nominalMagnification} | Magnification |
| {http://www.openmicroscopy.org/rdf/2016-06/ome_core/ContrastMethod} | Contrast |
| {http://www.openmicroscopy.org/rdf/2016-06/ome_core/Objective} | Microscope Objective |
| {http://rs.tdwg.org/dwc/terms/basisOfRecord} | Basis of Record |
| {http://rs.tdwg.org/dwc/terms/PreservedSpecimen} | Voucher specimen |
| {http://rs.tdwg.org/dwc/terms/EarliestDateCollected} | Collection Date |
| {http://rs.tdwg.org/dwc/terms/recordedBy} | Collector |
| {http://rs.tdwg.org/dwc/terms/identifiedBy} | Determined |
| {http://rs.tdwg.org/dwc/terms/geodeticDatum} | Geodetic datum |
| {http://www.ebi.ac.uk/efo/EFO_0005020} | Latitude |
| {http://www.ebi.ac.uk/efo/EFO_0005021} | Longitude |
| {http://rs.tdwg.org/dwc/terms/verbatimElevation} | Elevation |
| {http://rs.tdwg.org/dwc/terms/coordinatePrecision} | Precision |
| {http://purl.obolibrary.org/obo/OBI_0001048} | Camera |

## Data Records

Two separate data records were created in order to enable rapid use of the data in machine learning and biodiversity approaches.

1. The camera raw images (Canon CR3-format), the pre-processed images (16-bit TIFF-format), and the contextual meta-data were deposited in BioStudies under the identifier S-BIAD188 (https://www.ebi.ac.uk/biostudies/studies/S-BIAD188). The data record consists of a total of 223’989 individual raw image files partitioned into 48 species. The entire data record has a total size of approx. 12 TB.
2. The pre-processed and fully segmented and processed images along with meta-data were deposited in OME-TIFF and JPEG-format, respectively, in the Image Data Resource (IDR) repository under the identifier idr0134 (https://idr.openmicroscopy.org/search/?query=Name:134). The data record consists of a total of 4233 pre-processed and 905 fully processed imaged files. The data record has a total size of approx. 14 TB.

## Technical Validation

Biological validation of species identity and visible phenotypic attributes in the pre-processed images were performed by consulting the external experts Edwin Urmi, Heike Hofmann, Vadim Bakalin and Kristian Hassel. Photos of the herbarium specimen CM-30377 originating from North America (Table 1) show quite different properties when compared to the voucher specimen B-108428 originating from Northern Europe. Hence, the taxonomic status of the species *Scapania glaucocephala* is not yet fully clear^15^. Photos of CM-30377 may, thus, relate to the species *Scapania scapanioides* (C.Massal.) Grolle, which is listed in ^7^ as separate species occurring in Europe. Further, *S. brevicaulis* and *S. degenii* may comprise taxonomically identical species and additional research is needed to resolve their phylogenetic status. Images from this study can help the to clarify relationships of phenotypic attributes and the phylogenetic and taxonomic status of cryptic species.

To validate the technical quality of fused composite images, multi-focused image fusion methods were applied with different settings to the individual images in stacks. Composite images were manually inspected and the best image retained. Generally, classic Laplacian pyramid transform-based methods such as *Pyramid Maximum Contrast* implemented in the software Helicon Focus produce good results in complex cases with regard to intersecting objects, edges and many images, but these algorithms increase contrast and glare and they are prone to noise and artefacts and generally are considered less accurate regarding reproduction of microscopic objects^62–65^. The deterministic depth map-based method implemented in Helicon Focus first calculates depth maps from intermediary images based on the absolute difference in brightness of corresponding pixels in the source images and smoothed intermediary images and then generates the composite image from the source image pixels with indices differing from the indices in the smoothed depth map^66^. Whereas larger values for the parameter *radius* increase blur along edges, lower values can introduce artefacts, while the amount of blur along the transition between fused areas of individual images can be controlled with the parameter *smoothing*. While the depth map-based method generally produces accurate reproductions of microscopic objects, in some circumstances and especially with high magnifications, it can generate visible and large artefacts and blur around the edges (boundary regions) (Fig. 2). Recently, machine learning-based methods have been applied to focus-based image fusion tasks that may be superior to deterministic approaches^36^. Although there have been proposed some algorithms specifically for microscopic imaging, there is a considerable lack of usable implementations and a lack of microscopic training data for machine learning-based algorithms^36^.

**Figure 1:**
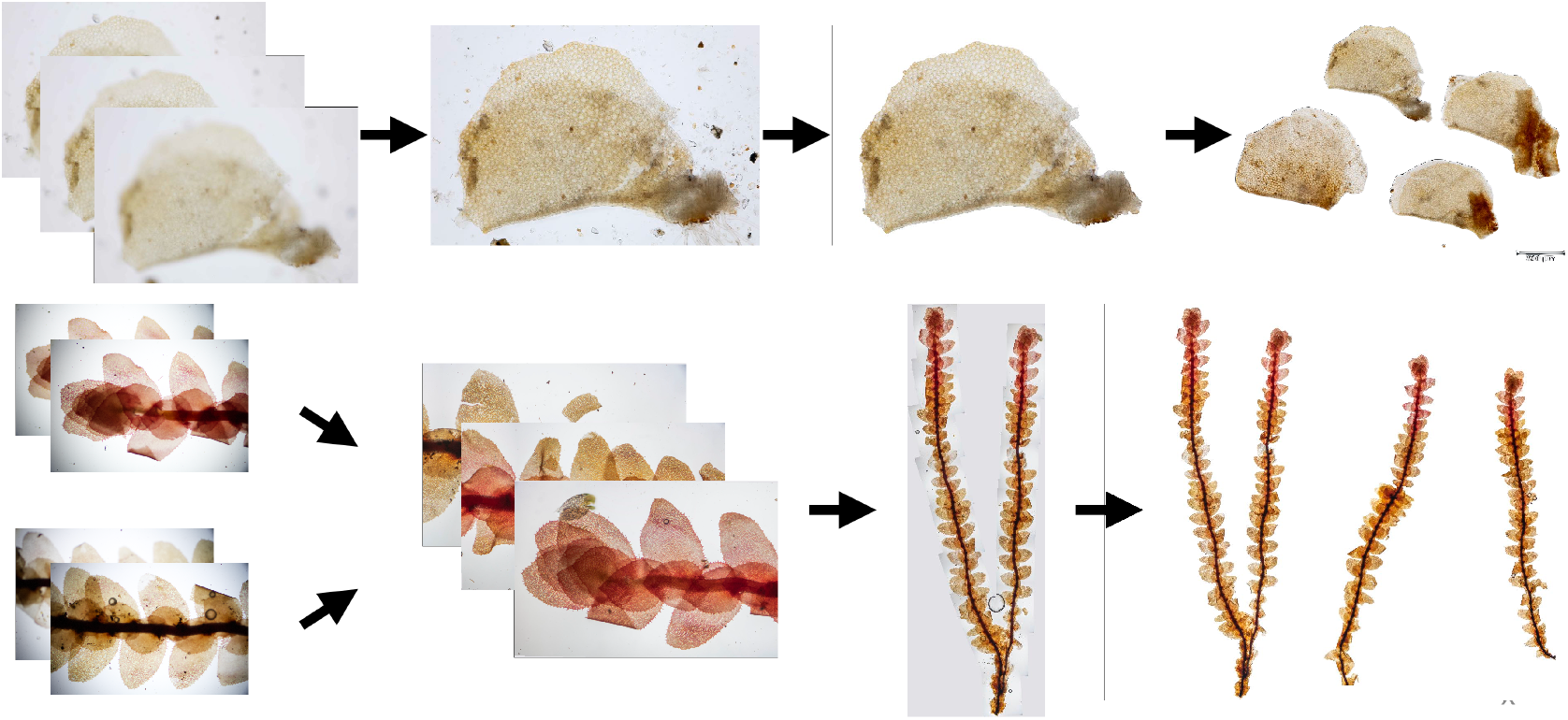
Two exemplary workflows used in this study to create segmented images. **(a)** Example of multi-focus image fusion where (1) several images of one object of a leaf lobe are fused into a (2) composite image. The leaf lobe as shown in the composite image is then (3) segmented and the background removed. Several leaf lobe objects are then put onto the (4) final image and a microscopic scale is applied. **(b)** Example of image stitching where (2) several fused images from (1) different segments of the same object are (3) stitched into a composite image with larger dimensions. Several of these stitched images are then put onto the (4) final image and a microscopic scale is applied.

**Figure 2:**
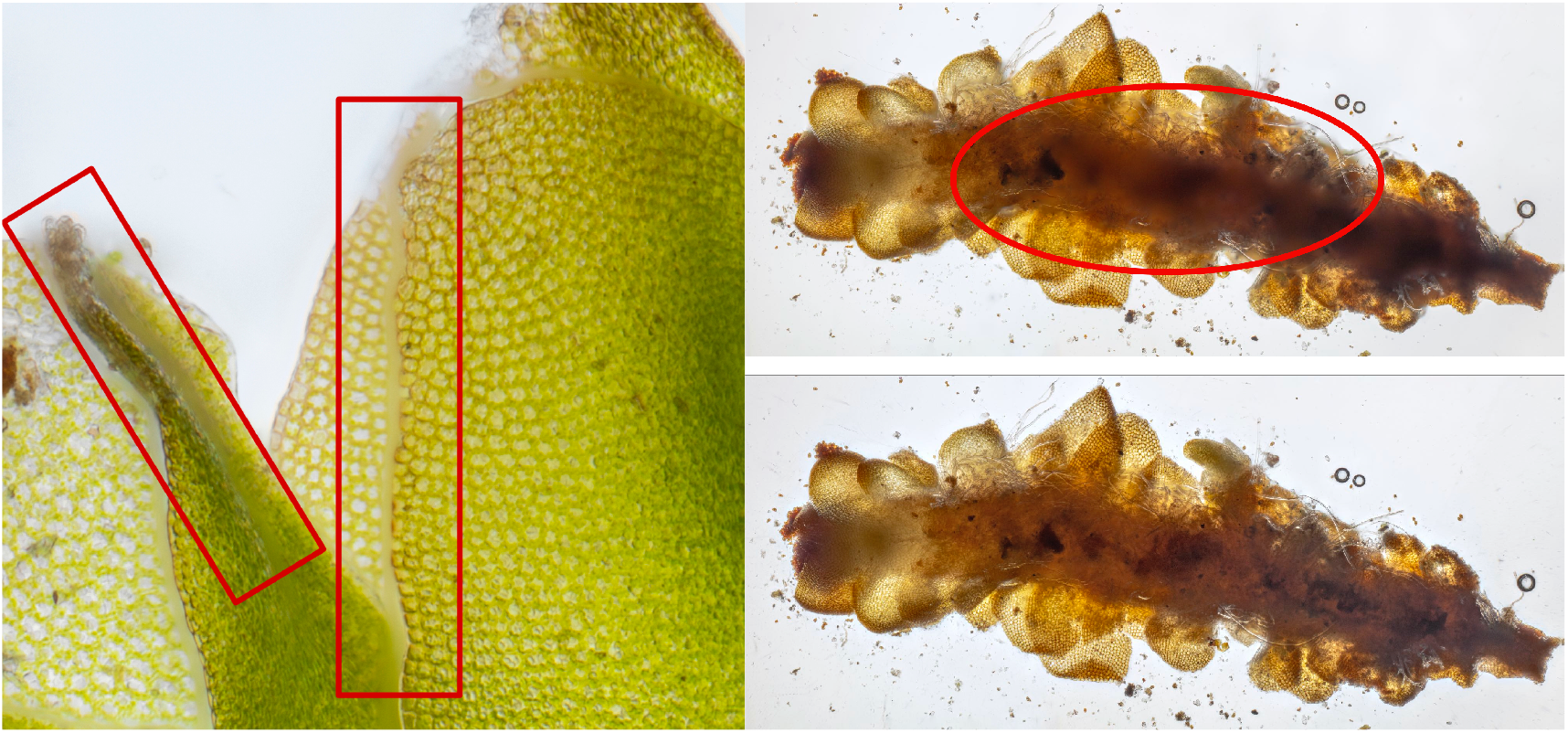
Deficiencies of multi-focus image fusion methods. Red circles and bars were drawn *post-hoc* with Affinity Photo to indicate the deficient regions in the images, thus, the regions where multi-focus image fusion methods can produce blur and artefacts in composite images of microscopic objects. **(a)** Crop of *IMG_1532-1621 Scapania cuspiduligera stature ventral side*. Visible blur along edges (boundary regions) of overlapping leaves (Parameter settings: *Method: Depth Map, Radius: 8, Smoothing: 4*). **(b)** *IMG_0107-0226 Scapania ligulifolia stature dorsal side* (Parameter settings: *Method: Depth Map, Radius: 32, Smoothing: 20*).

In order to facilitate the automatic processing of images, Python scripts have been written which are available as Open Source software in github^57,58^. These scripts use meta-data information to put individual images into image stacks to perform focus-based image fusion and image stitching tasks. Most of the work has been implemented manually. Scientific workflows allow to fully automate the entire process combining images with software tools utilising the machine-actionable information contained in the meta-data^42,48,67^. Meta-data has been validated using procedures described in ^20,45,46^ and standardised vocabularies were following the FAIR guiding principles^1^. When improved algorithms have been developed, the entire pipeline can be re-run resulting in improved segmented images without any further intervention. This data reuse and the rich documentation in meta-data will result in foster good scientific practices through source tracking and provenance.

## Usage Notes

Raw camera and pre-processed imaging data, the fully segmented and processed images, and the meta-data are available under the terms of the license Creative Commons CC BY 4.0. Open-Source software scripts and code^57,58^ are available under the terms of the BSD 3-Clause license.

## Code Availability

Software code and scripts used in this study are available as Open Source in github: https://github.com/korseby/create_image_stacks ^58^, https://github.com/korseby/scale_bar ^57^, https://github.com/korseby/bioimage_submission ^61^. Python scripts were tested under Python 3.7 and require the additional modules PIL, pandas, xml, csv, errno, sys, os, argparse, glob, pathlib and re. R scripts were tested under R 4.1.3 and require the additional packages parallel, foreach, and doMC. Shell scripts were tested using Bourne Again Shell (bash) 5.1.16.

## Acknowledgements

KP acknowledges the support of iDiv (funded by the German Research Foundation, DFG-FZT 118, 202548816) and Swissbryophytes. Fully segmented and processed images of 27 *Scapania* species occurring in Switzerland were originally produced for the website http://www.swissbryophytes.ch and are also available at the Zurich Open Repository Archive ^68–71,71–93^. We would also like to thank the external experts Edwin Urmi, Heike Hofmann, Vadim Bakalin and Kristian Hassel for providing voucher specimens and for performing validation of species identities on images of visible traits and especially Edwin Urmi for help with identification of fresh samples.

## Author contributions

KP performed the entire study. BKR supervised the study.

